# Augmenter of Liver Regeneration Regulates Cellular Iron Homeostasis by Modulating Mitochondrial Transport of ATP-Binding Cassette B8

**DOI:** 10.1101/2020.11.30.403295

**Authors:** Hsiang-Chun Chang, Jason S. Shapiro, Xinghang Jiang, Grant Senyei, Teruki Sato, Konrad T. Sawicki, Hossein Ardehali

## Abstract

Chronic loss of Augmenter of Liver Regeneration (ALR) results in mitochondrial myopathy with cataracts, however, the mechanism for this disorder remains unclear. Here, we demonstrate that loss of ALR, a principal component of the MIA40/ALR protein import pathway, results in impaired cytosolic Fe/S cluster biogenesis in mammalian cells. Mechanistically, MIA40/ALR facilitates the mitochondrial import of ATP binding cassette (ABC)-B8, an inner mitochondrial membrane protein required for cytoplasmic Fe/S cluster maturation, through physical interaction with ABCB8. Downregulation of ALR impairs mitochondrial ABCB8 import, reduces cytoplasmic Fe/S cluster maturation, and increases cellular iron through the iron regulatory protein-iron response element system. Our finding provides a mechanistic link between MIA40/ALR import machinery and cytosolic Fe/S cluster maturation through the mitochondrial import of ABCB8, and offers a potential explanation for the pathology seen in patients with ALR mutations.

## Introduction

Chronic loss of Augmenter of Liver Regeneration (ALR) results in mitochondrial myopathy and cataract, and combined respiratory chain deficiency (Di Fonzo, Ronchi et al. 2009, Calderwood, Holm et al. 2015, Nambot, Gavrilov et al. 2017). Several mutations in ALR have been described for this disorder. Some of these documented mutations result in premature stop codon, while the R194H mutation has been shown to affect the protein stability through increasing dissociation of cofactor FAD (Daithankar, Schaefer et al. 2010, Nambot, Gavrilov et al. 2017). Cellular studies demonstrated that R194H ALR is not functional, and mutant cells can be complemented by ectopic expression of wild type ALR (Di Fonzo, Ronchi et al. 2009). While patients with *ALR* mutations lack its protein expression, how reduced ALR protein levels are linked to human pathology remains to be determined.

ALR is a sulfhydryl oxidase with two isoforms– a full-length mitochondrial isoform and a shorter cytosolic isoform (Li, Wei et al. 2002). The cytosolic isoform of ALR, which is thought to function in hepatic regeneration (Li, Wei et al. 2002), lacks the first 80 amino acids, but retains the catalytic domain necessary to oxidize cysteines. The mitochondrial isoform of ALR localizes to the mitochondrial intermembrane space. Although the function of ALR is not entirely understood, it has been implicated in hepatic regeneration after injury (Polimeno, Pesetti et al. 2011), protection against hepatic and renal injury (Huang, Long et al. 2018, Liu, Xie et al. 2019), facilitation of cardiac development (Dabir, Hasson et al. 2013), and maintenance of embryonic stem cells (Todd, Gomathinayagam et al. 2010). Mice with liver-specific deletion of ALR from birth develop steatohepatitis and hepatocellular carcinoma, and have exacerbated hepatic injury when exposed to alcohol, while post-natal deletion of ALR in the liver induces oxidative stress and promotes hepatic steatosis (Gandhi, Chaillet et al. 2015, Kumar, Wang et al. 2016, Kumar, Rani et al. 2019). Previous studied demonstrated that ectopic expression of mitochondria-targeted human ALR rescues the cytoplasmic iron-sulfur (Fe/S) cluster maturation defects in *Erv1*-null yeast (Lange, Lisowsky et al. 2001), suggesting that ALR may be the functional homolog of Erv1p and play a role in Fe/S cluster biogenesis.

The synthesis of Fe/S clusters occurs in the mitochondria on the scaffold protein iron-sulfur cluster assembly enzyme (ISCU). Iron destined for Fe/S cluster synthesis is imported into mitochondria by mitoferrin 1/2 and is delivered to ISCU by frataxin. The fully-assembled Fe/S clusters are then transferred to Fe/S cluster-containing proteins. Because several cytosolic proteins also require Fe/S clusters for their function, another set of cytosolic proteins, collectively named as the cytosolic iron/sulfur cluster assembly (CIA) machinery, facilitates the incorporation of Fe/S clusters into cytosolic proteins (Lill, Dutkiewicz et al. 2006). However, the identity of the molecule that supplies CIA machinery and how it gets out of mitochondria has not been identified. Proper cytosolic Fe/S cluster maturation is critical for cell health. Deletion of ABCB8 in the heart, which leads to cytosolic Fe/S cluster deficiency, results in spontaneous cardiomyopathy (Ichikawa, Bayeva et al. 2012). Conditional deletion of *Iop1*, one component of CIA machinery, in mice results in complete lethality, and deletion of this gene in mouse embryonic fibroblasts lead to cellular iron accumulation and rapid cell death (Song and Lee 2011). Previous studies in yeast have implicated three essential players for cytoplasmic Fe/S cluster biogenesis – Atm1p, Erv1p and glutathione (Lill, Dutkiewicz et al. 2006). Additionally, Erv1 has also been shown to mediate cytosolic Fe/S cluster biogenesis in the parasitic trypanosome *T. brucei* (Haindrich, Boudova et al. 2017).

To support adequate Fe/S cluster biogenesis, iron first enters the cells through transferrin-receptor mediated uptake. It can subsequently be stored in the cytoplasm or incorporated into heme and iron-sulfur (Fe/S) clusters in the mitochondria (Hentze, Muckenthaler et al. 2010, Ye and Rouault 2010). While adequate supply of iron is required for normal cellular function, excess iron can catalyze the formation of reactive oxygen species. Therefore, the levels of cellular iron are tightly regulated. The iron regulatory protein (IRP)-iron response element (IRE) system is the master regulator of cellular iron in mammals. There are two IRPs in mammalian systems, IRP1 and IRP2. In the iron-replete state, IRP1 acquires an Fe/S cluster and functions as a cytosolic aconitase, while IRP2 is proteolytically degraded in an iron- and oxygen-dependent process. In the iron-starved state, IRP1 loses its Fe/S cluster and becomes activated, and IRP2 is stabilized. Both proteins then bind to IREs within the 5’- and 3’-untranslated regions (UTRs) of mRNA and post-transcriptionally regulate gene expression (De Domenico, McVey Ward et al. 2008, Hentze, Muckenthaler et al. 2010, Anderson, Shen et al. 2012).

Erv1p, together with Mia40, is part of the oxidative folding machinery required for the import of a subset of mitochondrial proteins. Mia40 oxidizes the cysteine residues within imported proteins, and the reduced Mia40 is re-oxidized by Erv1p, effectively recycling Mia40 for further use (Müller, Milenkovic et al. 2008). A similar electron transfer process has also been demonstrated using recombinant human MIA40 and ALR proteins (Banci, Bertini et al. 2011). A subset of proteins imported by the Mia40/Erv1p machinery, including Tim 13 and Tim 22, are involved in the mitochondrial import of other proteins (Müller, Milenkovic et al. 2008, Wrobel, Trojanowska et al. 2013). Nevertheless, the functional significance of these isoforms on cytosolic Fe/S cluster maturation and cellular iron regulation in mammalian systems remains unclear.

In this paper, we demonstrate that cytoplasmic Fe/S cluster maturation depends on a functional mitochondrial protein import system consisting of ALR and MIA40. Mitochondrial ALR facilitates the mitochondrial import of ABCB8, a protein needed for the maturation of cytosolic Fe/S proteins. Thus, a reduction in ALR and eventual reduction in the cytosolic Fe/S clusters leads to activation of IRP1 and subsequent cellular iron accumulation.

## Results

### Downregulation of *Alr* results in cytosolic Fe/S maturation defects

Previous yeast studies demonstrated that loss of Erv1p resulted in cytosolic Fe/S cluster deficiency, which was rescued with forced overexpression of human ALR (Lange, Lisowsky et al. 2001). Although the defect is restricted to cytosolic protein, humans with chronic loss of ALR demonstrates mitochondrial myopathy with respiratory chain defect (Di Fonzo, Ronchi et al. 2009, Calderwood, Holm et al. 2015, Nambot, Gavrilov et al. 2017). To elucidate the role of ALR in cytosolic Fe/S cluster maturation and mitochondrial function in mammalian cells, we first downregulated *Alr* in mouse embryonic fibroblasts (MEFs) using pooled siRNA and demonstrated a significant reduction of *Alr* at both mRNA and protein levels (***Figure 1A and B***). As the quantity of cytosolic Fe/S clusters cannot be measured directly, the protein levels of glutamine PRPP amidotransferases (GPAT), a cytosolic protein whose maturation requires Fe/S clusters, and the activities of xanthine oxidase (XO) and cytosolic aconitase, both of which use Fe/S clusters as cofactors, were used as surrogate markers for cytosolic Fe/S cluster maturation. Downregulation of *Alr* decreased GPAT protein levels (***Figure 1C***), and reduced XO (***Figure 1D***) and cytosolic aconitase activity (***Figure 1E***). We also measured the activity of the Fe/S cluster-containing mitochondrial complexes I and II, and found no change in complex I and II activity with *Alr* downregulation (***Figure 1F and G***). Our results demonstrated that acute reduction in ALR impairs specifically cytosolic Fe/S cluster maturation in mammalian cells similar to the previously published yeast studies.

**Figure 1.**
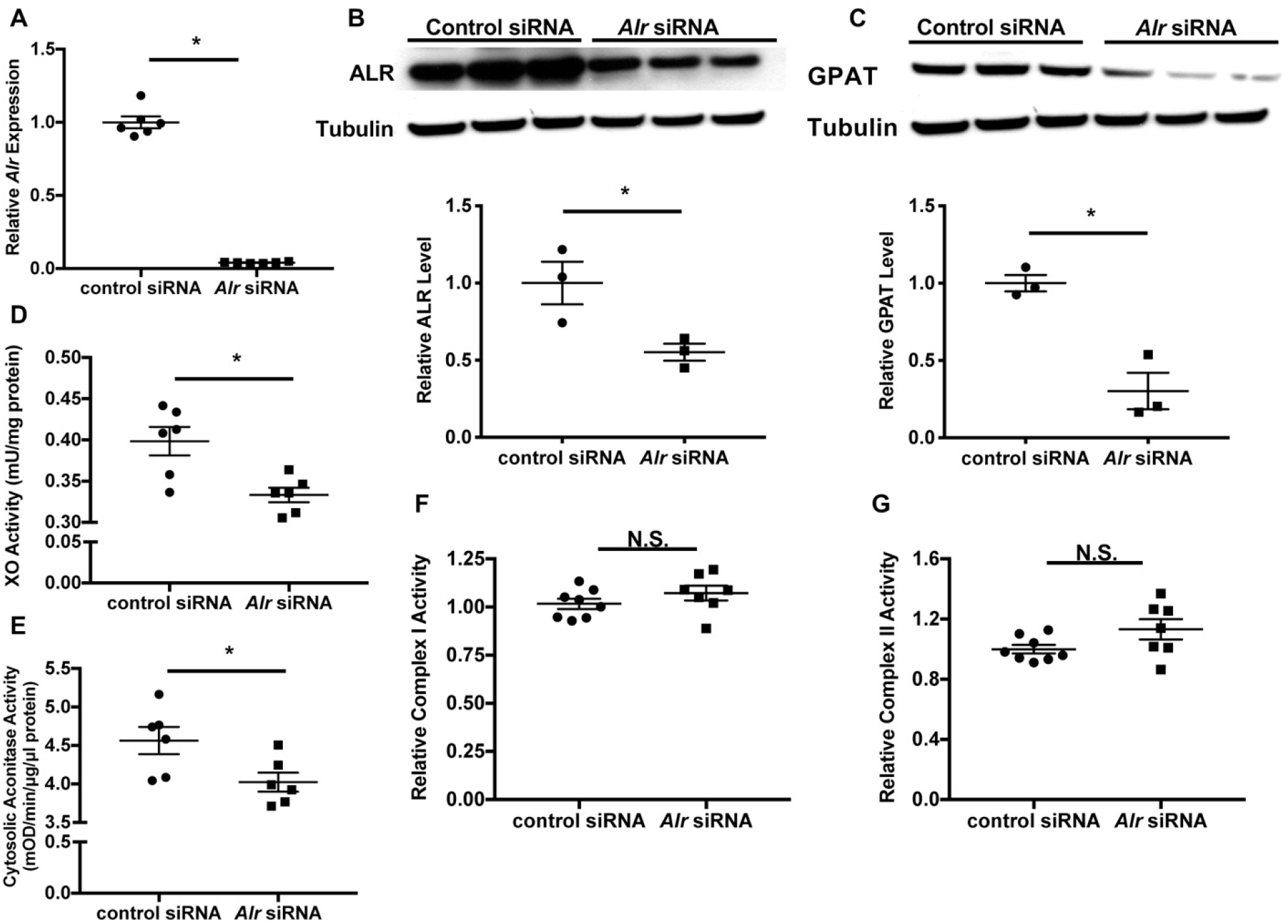
Downregulation of *Alr* results in cytoplasmic Fe/S cluster maturation defects. (**A**) mRNA (n=6) and (**B**) protein (n=3) levels of *Alr* in wild type (WT) MEFs with or without *Alr* downregulation. GPAT protein levels (n=3, **C**), xanthine oxidase (XO, n=6, **D**) and cytosolic aconitase (n=6, **E**) activities in cells treated with *Alr* siRNA. Mitochondrial complex I (n=7-8, **F**) and complex II activity (n=7-8, **G**) in WT MEFs with *Alr* downregulation. Quantification of western blotting images is shown in the same panel. Data are presented as mean ± SEM. * P<0.05 by ANOVA. N.S.=not significant.

### Downregulation of *Alr* increases cellular iron through activation of the IRP-IRE system

We next investigated changes in cellular iron with *Alr* downregulation. Transferrin receptor-1 (*Tfrc*) mRNA levels were increased in wild type MEFs with *Alr* downregulation (***Figure 2A***). Consistent with changes in *Tfrc* expression, we also observed an increase in transferrin-mediated iron uptake in cells treated with *Alr* siRNA (***Figure 2B***). The increased iron uptake resulted in higher steady state levels of iron in both the cytoplasm and mitochondria (***Figure 2C***), similar to other genetic models with cytosolic Fe/S cluster maturation defects (Sato, Torimoto et al. 2011, Ichikawa, Bayeva et al. 2012). Since nontransferrin-bound iron is not present in normal culturing conditions, these results suggest that downregulation of *Alr* increases cellular iron, likely through upregulation of *Tfrc*.

**Figure 2.**
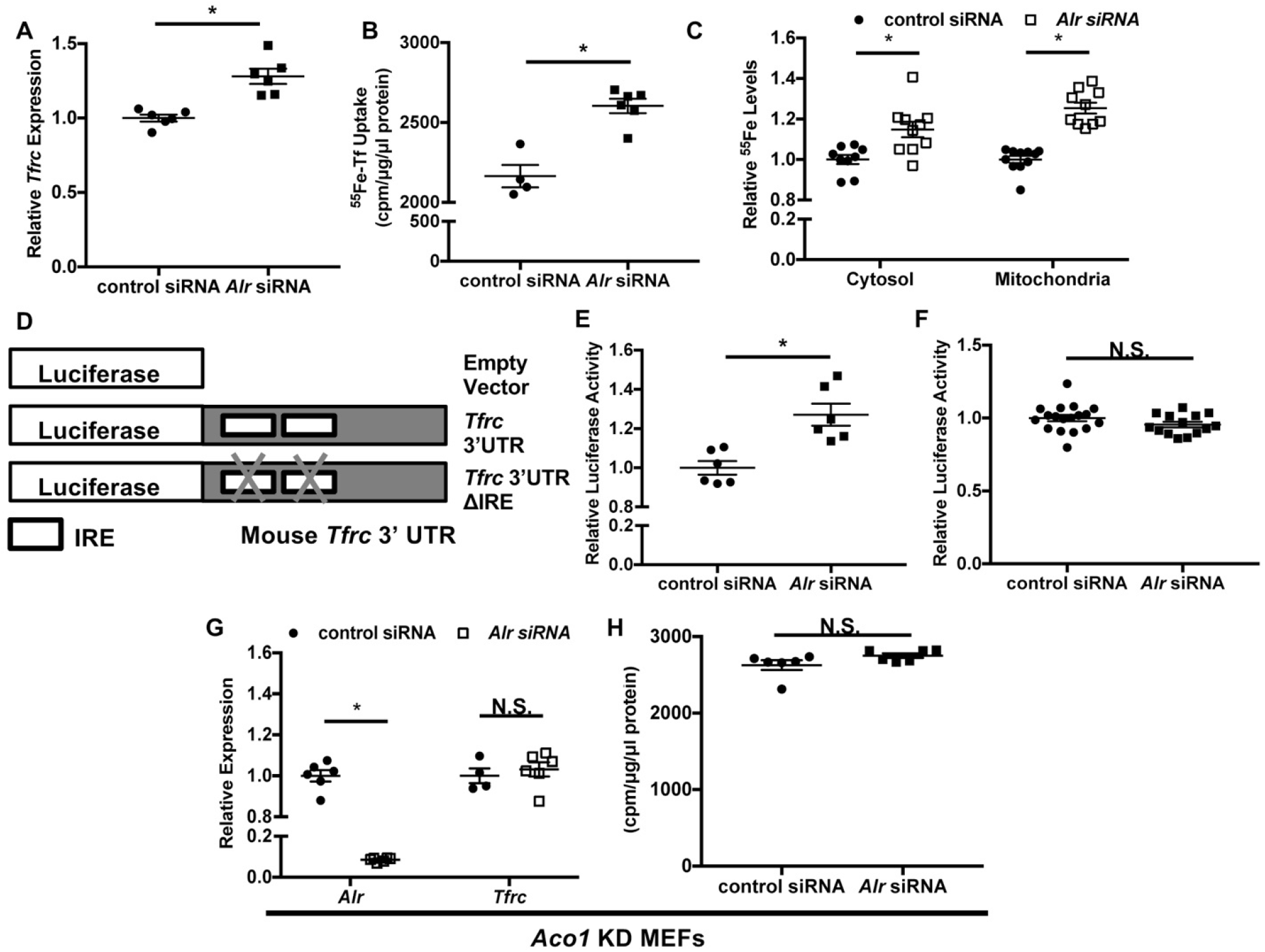
Downregulation of ALR increases *Tfrc* levels and cellular iron content through IRP1. *Tfrc* mRNA levels (**A**) and transferrin-dependent iron uptake (**B**) in WT MEFs with *Alr* downregulation (n=5-6). (**C**) Steady-state ^55^Fe levels in cytosolic and mitochondrial fraction from cells treated with *Alr* siRNA (n=6). (**D**) Schematic representation of *Tfrc* 3’UTR reporter constructs. (**E**) Full length *Tfrc* 3’UTR reporter activities were measured in WT MEFs with ALR downregulation (n=6). (**F**) IRE-deleted *Tfrc* 3’UTR reporter activities in WT MEFs with ALR downregulation (n=14-18). (**G**) *Alr* and *Tfrc* mRNA levels in *Aco1* KD MEFs treated with *Alr* siRNA (n=4-6). (**H**) TFRC-mediated ^55^Fe uptake in *Aco1* KD MEFs treated with *Alr* siRNA (n=6). Data are presented as mean ± SEM. * P<0.05 by ANOVA. N.S.=not significant.

Both transcriptional and post-transcriptional regulation have been shown to alter *Tfrc* mRNA levels (Tacchini, Gammella et al. 2008, Anderson, Shen et al. 2012, Bayeva, Chang et al. 2013). We therefore next studied the mechanism for the upregulation of *Tfrc* by *Alr* knockdown. We first generated a luciferase reporter construct harboring 700 base pairs of the *Tfrc* promoter upstream of the *Tfrc* transcription start site (***Figure 2-figure supplement 1A***). Downregulation of *Alr* did not change luciferase activity of this construct (***Figure 2-figure supplement 1B***), suggesting that ALR does not regulate *Tfrc* expression at the transcriptional level. We next made a construct containing the full length *Tfrc* 3’-UTR downstream of luciferase cDNA (***Figure 2D***). Luciferase activity of this reporter increased with iron chelation (***Figure 2-figure supplement 1C***), and with *Alr* downregulation (***Figure 2E***), suggesting that ALR post-transcriptionally regulates *Tfrc* through stabilization of *Tfrc* mRNA.

Since *Tfrc* is modulated at the mRNA level by IRPs, we then studied the role of IRP1/2 in ALR-mediated changes in *Tfrc*. We first made a luciferase construct with all IRE sites removed from the *Tfrc* 3’-UTR (***Figure 2D***). As expected, this construct displayed no responsiveness to iron chelation (***Figure 2-figure supplement 1D***). Furthermore, downregulation of *Alr* had no effect on the luciferase activity of the IRE-deleted reporter construct (***Figure 2F***), suggesting that activation of the IRP/IRE system is required for the effects of ALR on cellular iron homeostasis.

We then assessed the role of IRP2 by treating *Ireb2* (encoding IRP2) knockout (KO) MEFs with *Alr* siRNA. *Alr* knockdown in these cells resulted in a significant reduction in ALR protein levels (***Figure 2-figure supplement 1E***), and a significant increase in *Tfrc* mRNA levels and transferrin-mediated iron uptake (***Figure 2-figure supplement 1 F and G***), suggesting that the regulation of *Tfrc* by ALR is independent of IRP2. Thus, we focused our studies on Irp1. We first created a stable MEF line expressing *Aco1* (encoding IRP1) shRNA, but we only achieved 40% downregulation of *Aco1* at the mRNA level (***Figure 2-figure supplement 1H***). To more effectively downregulate *Aco1* expression, we treated cells expressing *Aco1* shRNA with siRNA against *Aco1* (referred to as *Aco1* KD MEFs hereafter) and achieved a significant decrease in *Aco1* transcript levels (***Figure 2-figure supplement 1I***). Although treatment of *Aco1* KD MEFs with *Alr* siRNA significantly reduced *Alr* mRNA (***Figure 2G***), *Tfrc* mRNA levels and TFRC-mediated iron uptake were not increased (***Figure 2G and H***). Since IRP1 is activated upon losing its mature Fe/S cluster, our results indicate that *Alr* downregulation impairs Fe/S cluster maturation and subsequently promotes increases in *Tfrc* mRNA and cellular iron levels in an IRP1-dependent manner.

### The mitochondrial isoform of ALR regulates cellular iron and cytosolic Fe/S cluster maturation

The ALR gene encodes two isoforms: a full length mitochondrial isoform, and a shorter cytoplasmic isoform lacking the first 80 amino acids (Li, Wei et al. 2002). To determine which isoform of ALR regulates cellular iron and the maturation of cytosolic Fe/S clusters, we generated lentiviral constructs containing the protein coding regions of the two isoforms of human ALR, but lacking 3’-UTR sequences. Using these constructs, we overexpressed these two isoforms in MEFs and confirmed the localization of each isoform to their respective cellular compartments (***Figure 3-figure supplement 1A and B***).

We then knocked-down endogenous *Alr* with siRNA targeting the 3’-UTR of mouse *Alr* mRNA, followed by lentiviral overexpression of either isoform using the aforementioned virus. This approach allowed us to selectively overexpress individual isoforms of ALR in the absence of the endogenous ALR protein. Quantitative RT-PCR using species-specific primer sets confirmed that only endogenous, but not exogenous, *Alr* was downregulated by siRNA (***Figure 3-figure supplement 1C and D***). Overexpression of the full-length (mitochondrial) but not the short (cytosolic) isoform of ALR rescued the increase in *Tfrc* mRNA levels and transferrin-dependent iron uptake (***Figure 3A and B***). In addition, the cytosolic Fe/S cluster maturation defects, including reduced cytosolic aconitase and XO activity, and decreased GPAT protein levels, were all reversed by the re-expression of the mitochondrial but not the cytosolic isoform of ALR (***Figure 3 C-F***). These observations indicate that the mitochondrial ALR isoform regulates cytosolic Fe/S cluster maturation, which in turn affects IRP1 activity and cellular iron levels.

**Figure 3.**
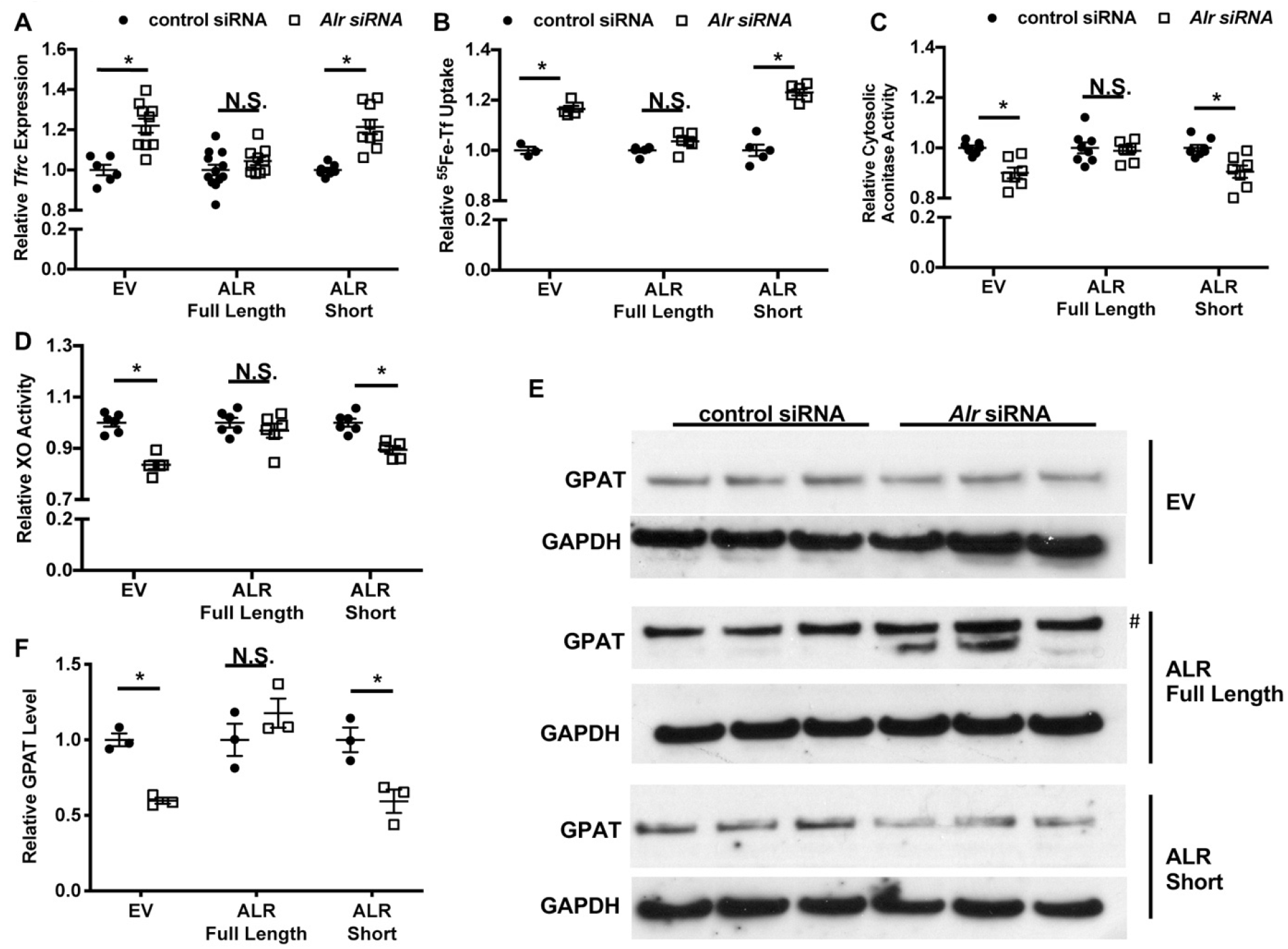
Overexpression of mitochondrial but not cytosolic isoform of ALR rescues the iron-sulfur cluster maturation defects from ALR downregulation. *Tfrc* mRNA levels (n=6, **A**) and transferrin-dependent iron uptake (n=4-6, **B**) were measured in WT MEFs with endogenous *Alr* downregulation and concurrent overexpression of different ALR isoforms. Cytosolic aconitase (n=7-8, **C**) and XO activities (n=5-6, **D**), and GPAT protein levels (n=3, **E**) are measured in cells with downregulation of endogenous ALR and overexpression of various ALR isoforms. **#** indicate the band correspond to GPAT protein. (**F**) Quantification of panel **E**. EV=empty vector. ALR short=cytosolic isoform of ALR lacking the first 80 amino acids. Data are presented as mean ± SEM. * P<0.05 by ANOVA with *post hoc* Tukey comparison. N.S.=not significant.

### ALR deficiency impairs mitochondrial transport of ABCB8

Our results thus far indicate that mitochondrial ALR is required for cytosolic Fe/S cluster maturation; however, the mechanism by which ALR carries out this function is not clear. ALR downregulation results in Fe/S cluster maturation defects in the cytoplasm but not in the mitochondria, and this phenotype bears striking similarity to cells with deficiency in ABCB7 or ABCB8 (Pondarré, Antiochos et al. 2006, Ichikawa, Bayeva et al. 2012), two mitochondrial proteins involved in the maturation of cytoplasmic Fe/S clusters. We therefore hypothesized that ALR influences cytosolic Fe/S cluster maturation by modulating the levels of ABCB7 and/or ABCB8. Since mitochondrial functional studies would require large quantities of mitochondria not attainable from MEFs, we used HEK293 cells for the next set of studies. First, we showed that siRNA treatment resulted in a significant decrease in *ALR* mRNA and protein levels in HEK293 cells (***Figure 4-figure supplement 1A and B***). HEK293 cells with *ALR* downregulation also demonstrated cytosolic Fe/S cluster maturation defects as measured by GPAT protein levels (***Figure 4-figure supplement 1C and D***). However, the mRNA levels of *ABCB7* and *ABCB8* (***Figure 4A***), and total cellular ABCB7 and ABCB8 protein levels (***Figure 4B and C***) did not change with ALR downregulation.

**Figure 4.**
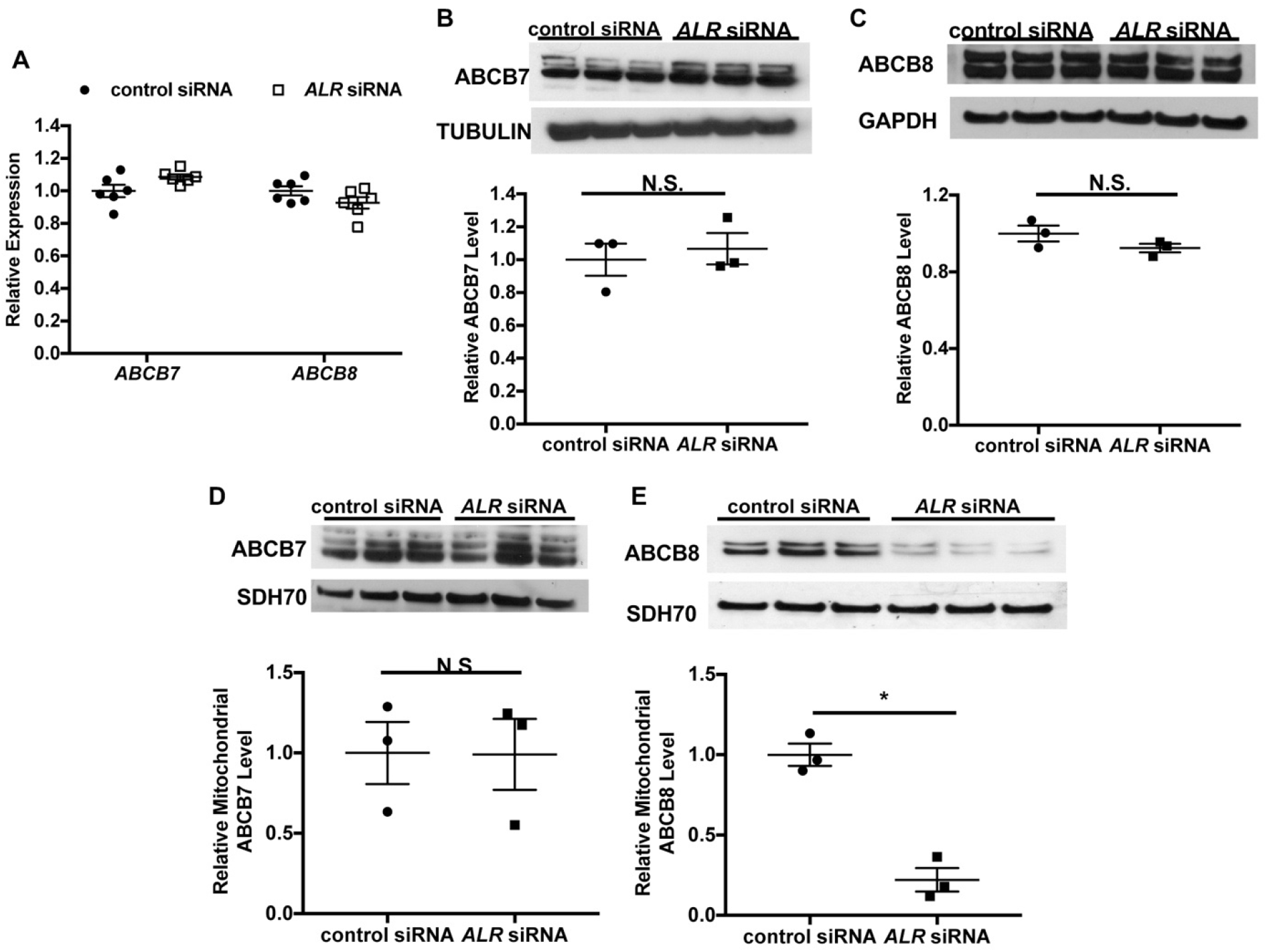
Downregulation of *ALR* in HEK293 cells reduces mitochondrial ABCB8 protein level. (**A**) mRNA levels of ABCB7 and ABCB8 in HEK293 cells with *ALR* downregulation (n=6). Total cellular levels of ABCB7 (**B**) and ABCB8 (**C**) in HEK293 cells with *ALR* downregulation. Mitochondrial levels of ABCB7 (**D**) and ABCB8 (**E**) in HEK293 cells with *ALR* downregulation. Densitometry analysis is shown together with representative images. n=3 for panels **B-E**. Data are presented as mean ± SEM. * P<0.05 by ANOVA. N.S.=not significant.

As ALR is involved in mitochondrial protein import, we next examined the amount of ABCB7 and ABCB8 protein contained within the mitochondrial fraction from cells with *ALR* downregulation. *ALR* downregulation did not change the mitochondrial level of ABCB7 (***Figure 4D***), but caused a significant reduction in mitochondrial ABCB8 (***Figure 4E***). Additionally, ABCB7 overexpression did not rescue the increase in *Tfrc* expression in ALR-downregulated MEFs (***Figure 4-figure supplement 1E***), demonstrating lack of functional overlap between ABCB7 and ABCB8. These results suggest that ALR may be required for the mitochondrial import of ABCB8 but not ABCB7.

To determine whether ALR influences the mitochondrial transport of ABCB8, we generated a doxycycline (DOXY)-inducible expression system that allows for monitoring of protein accumulation after DOXY treatment. HEK293 cells stably expressing reverse tetracycline transactivator (rtTA3) were transfected with a tetracycline responsive bicistronic construct containing GFP and a 6X His-tagged gene of interest. This system results in mitochondrial protein accumulation in a time-dependent manner in response to DOXY treatment (***Figure 5-figure supplement 1A and B***). To correct for differences in transfection efficiency, the mitochondrial level of the His-tagged protein was normalized to the cytosolic level of GFP.

Mammalian MIA40 does not contain a presequence needed for mitochondrial targeting. Instead, its folding and retention in the mitochondria requires a functional oxidative folding system consisting of MIA40 and ALR, which are already localized to the mitochondrial intermembrane space (Sztolsztener, Brewinska et al. 2013). We first tested the function of the existing import system by showing that the mitochondrial import of newly synthesized MIA40 is decreased with *ALR* downregulation (***Figure 5A***). As a negative control, we examined the mitochondrial import of an eGFP fusion protein containing the first 69 amino acids of ATP synthase subunit 9 (SU9), which serves as a mitochondrial localization sequence and mediates mitochondrial import through a membrane potential-dependent but MIA40/ALR-independent mechanism (Wrobel, Trojanowska et al. 2013). This construct is subsequently referred to as SU9-eGFP. As expected, mitochondrial import of the His-tagged SU9-eGFP did not change with *ALR* downregulation (***Figure 5-figure supplement 1C***). We then studied the effects of ALR downregulation on the mitochondrial import of ABCB7 and ABCB8. *ALR* downregulation resulted in reduced mitochondrial import of ABCB8 (***Figure 5B***), but not ABCB7 (***Figure 5C***). These findings suggest that *ALR* impairs the mitochondrial import of ABCB8, which in turn affects cytosolic Fe/S cluster maturation.

**Figure 5.**
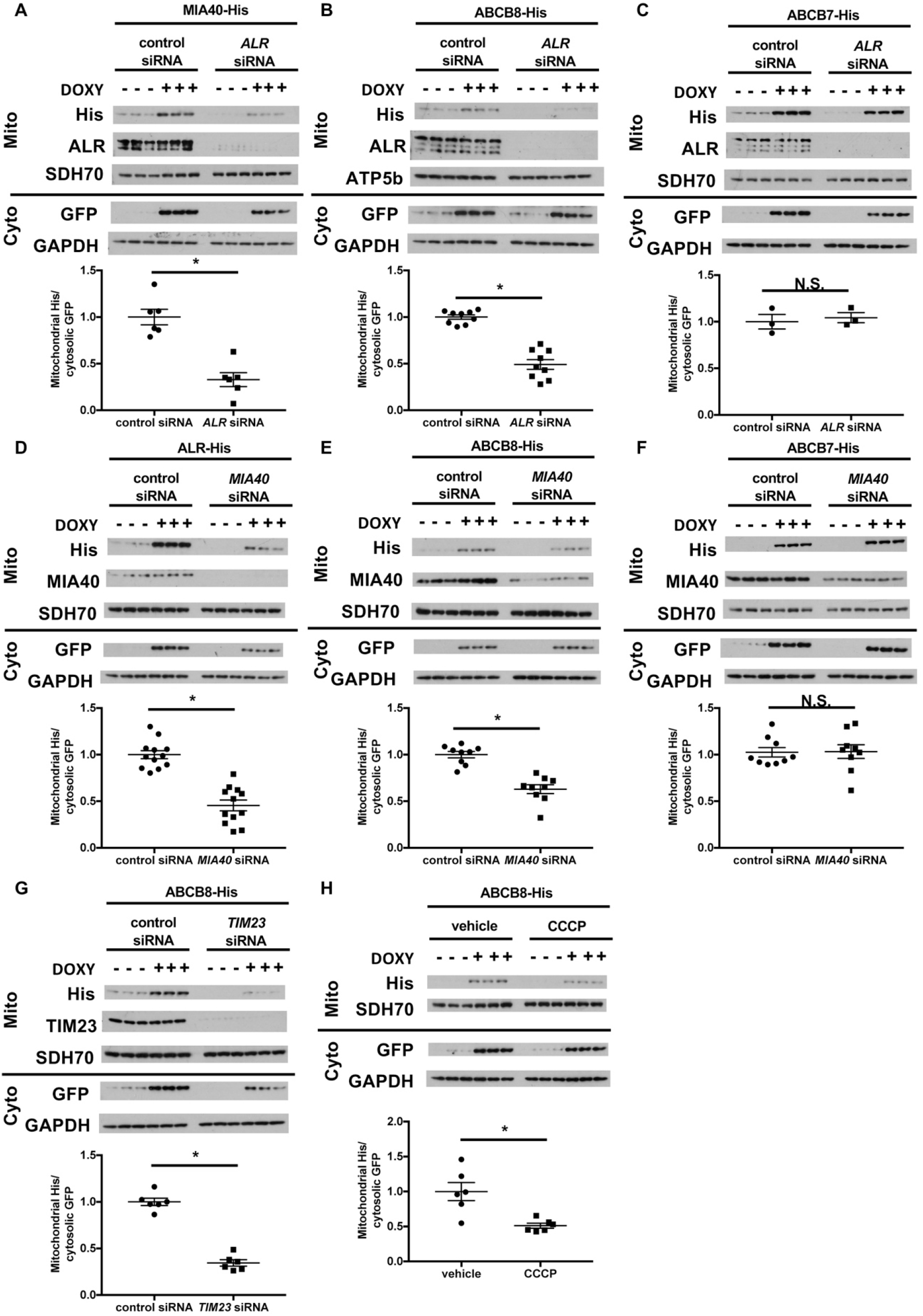
Mitochondrial import of ABCB8 requires ALR, MIA40, TIM23 and mitochondrial membrane potential. Imported levels of MIA40-His (n=6, **A**), whose mitochondrial translocation depends on Mia40/ALR protein transport pathway, ABCB8-His (n=9, **B**), and ABCB7-His (n=3, **C**) in cells treated with ALR siRNA with and without DOXY treatment for 6 hours. Imported levels of ALR-His (n=12, **D**), ABCB8-His (n=9, **E**), and ABCB7-His (n=9, **F**) in cells with or without *MIA40* downregulation with and without DOXY treatment for 6 hours. (**G**) Imported levels of ABCB8-His in cells with or without *MIA40* downregulation with and without DOXY treatment for 6 hours (n=6). (**H**) Imported levels of ABCB8-His in cells treated with CCCP and/or DOXY (n=6). Representative western blotting results and densitometry analyses are shown in each panel. Mito=mitochondrial fraction. Cyto=cytosolic fraction. Data are presented as mean ± SEM. * P<0.05 by ANOVA. N.S.=not significant.

### ABCB8 interacts with MIA40/ALR protein import machinery prior to its transport by the TIM23 complex

While the direct involvement of ALR in mitochondrial protein transport has not been described, ALR is known to play a critical role in the MIA40-mediated protein import pathway by re-oxidizing MIA40 (Kallergi, Andreadaki et al. 2012). We therefore tested whether downregulation of MIA40 recapitulated the defects of mitochondrial protein transport observed with ALR downregulation. We first confirmed that *MIA40* downregulation in HEK293 cells resulted in increased *TFRC* expression (***Figure 5-figure supplement 1D and E***), similar to what we observed with *ALR* downregulation (***Figure 2A***). In cells with *MIA40* downregulation, we observed a decrease in mitochondrial import of ALR, a known substrate of MIA40 (Kallergi, Andreadaki et al. 2012), but not SU9-eGFP (***Figure 5D and Figure 5-figure supplement 1F***). Similar to our finding with *ALR* downregulation, *MIA40* downregulation impaired mitochondrial import of ABCB8 but not ABCB7 (***Figure 5E and F***). These observations collectively indicate that mitochondrial import of ABCB8 requires both MIA40 and ALR.

Similar to many other inner mitochondrial membrane proteins transported by the TIM23 complex (Jensen and Dunn 2002), ABCB8 also contains a cleavable N-terminal targeting sequence. We therefore tested whether it is also transported by the TIM23 complex. Downregulation of TIM23 (an integral protein of the TIM23 complex) resulted in decreased mitochondrial import of both SU9-eGFP (***Figure 5-figure supplement 1G,*** positive control) and ABCB8 (Fig. 5G), while the mitochondrial import of MIA40 (the negative control) was not affected (***Figure 5-figure supplement 1H***). Additionally, dissipation of mitochondrial membrane potential, a required factor for TIM23 mediated protein transport (Jensen and Dunn 2002), decreased mitochondrial transport of SU9eGFP and ABCB8 (***Figure 5H and Figure 5-figure supplement 1I***). These results collectively support a model in which, after passing through the outer mitochondrial membrane, ABCB8 peptide interacts with the MIA40 protein import complex prior to being transported to the inner mitochondrial membrane by the TIM23 complex. A similar mechanism has been described for the yeast mitochondrial ribosomal protein Mrp10 (Longen, Woellhaf et al. 2014).

To test whether ABCB8 directly interacts with MIA40, we overexpressed C-terminal His-tagged MIA40 in HEK293 cells and loaded the isolated mitochondrial lysate onto a Ni-NTA column to pull down MIA40 and its interacting proteins. Endogenous ABCB8 copurified with MIA40 (***Figure 6A***), suggesting that ABCB8 interacts with MIA40 during its translocation into the mitochondria. Mutation of either or both of the two cysteine residues in MIA40, which have been shown to be required for substrate interaction (Peleh, Cordat et al. 2016), abolished the interaction between MIA40 and ABCB8 (***Figure 6B***).

**Figure 6.**
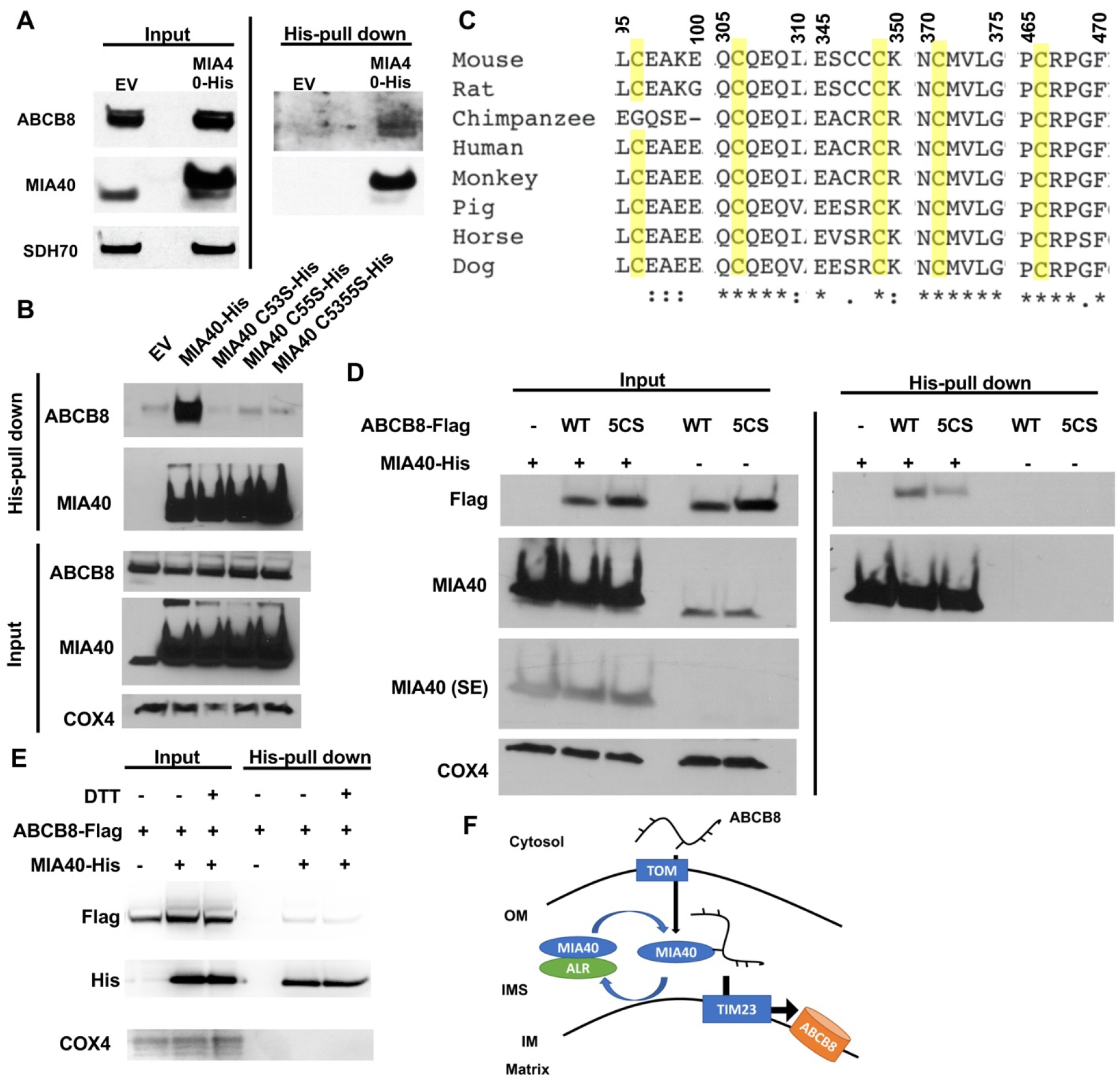
ABCB8 interacts with Mia40 during its transport into mitochondria. (**A**) Representative western blot image of mitochondrial lysates and His-tag pulldown fraction from cells overexpressing His-tagged MIA40 or empty vector. (**B**) Co-immunoprecipitation of His-tagged MIA40 and endogenous ABCB8 in mitochondrial fraction from cells overexpressing indicated constructs. (**C**) Clustal Omega alignment of ABCB8 protein sequences from higher vertebrate showing the location of five conserved cysteines (highlighted in yellow). (**D**) Co-immunoprecipitation of His-tagged MIA40 and Flag-tagged ABCB8 in mitochondrial fraction from cells overexpressing indicated constructs. (**E**) Co-immunoprecipiration of His-tagged MIA40 and Flag-tagged ABCB8 in mitochondrial fraction in the presence and absence of 100mM DTT. (**F**) Schematic representation of proposed model in which ABCB8 peptide interact with the MIA40/ALR protein import machinery prior to insertion into inner mitochondrial membrane through TIM23 complex. EV=empty vector. 5CS=ABCB8-Flag construct with mutations of five conserved cysteines to serine. SE=short exposure. OM=outer mitochondrial membrane. IMS=intermembrane space. IM=inner mitochondrial membrane.

Since MIA40 recognizes cysteine residues on target proteins to facilitate their mitochondrial import (Sideris, Petrakis et al. 2009), we performed alignment of ABCB8 protein sequences in vertebrates, and identified five highly conserved cysteine residues (***Figure 6C***). While mutating each of these residues had minimal effect on the interaction between ABCB8 and Mia40 (***Figure 6-figure supplement 1A***), mutation of all five cysteines to serines (5CS-ABCB8) greatly diminished the interaction of ABCB8 to MIA40 (***Figure 6D***). To further support this model, we perform pulldown experiment of wild type MIA40 and ABCB8 in the presence of 100mM dithiothreitol (DTT). DTT greatly diminished the interaction between MIA40 and ABCB8 (***Figure 6E***), similar to the 5CS mutant. The five conserved cysteine residues are not conserved between ABCB7 and ABCB8 (***Figure 6-figure supplement 1B***), thereby explaining the differential dependency of MIA40/ALR protein import system on mitochondrial import of these two proteins. Thus, these 5 cysteine residues are collectively needed for recognition by MIA40, and the absence of one is not sufficient to disrupt the interaction between ABCB8 and MIA40. Collectively, these observations suggest that MIA40 and ABCB8 interact through disulfide bond formation.

## DISCUSSION

Mutations in ALR has been linked to mitochondrial myopathy (Di Fonzo, Ronchi et al. 2009, Calderwood, Holm et al. 2015, Nambot, Gavrilov et al. 2017), but how dysfunctional ALR drives the disease phenotype remained to be determined. Previous studies hint that ALR may function similar to its yeast homolog Erv1p and involve in cytosolic Fe/S cluster maturation (Lange, Lisowsky et al. 2001). We hereby demonstrated that through its role in mitochondrial intermembrane space, ALR is critical for cytosolic Fe/S cluster biogenesis. The iron overload phenotype of cells with ALR deletion is reminiscent of loss of components of the CIA machinery (Song and Lee 2011) or other mitochondrial protein required for cytosolic Fe/S cluster biogenesis (Ichikawa, Bayeva et al. 2012). Importantly, we discover that downregulation of ALR prevents mitochondrial import of ABCB8, and overexpression of mutant ALR does not rescue the iron overload phenotype and impaired ABCB8 import. Our paper thus elucidated the molecular link between ALR and cytosolic Fe/S cluster maturation in mammalian system.

Patients with mutations in *ALR* that result in reduced ALR protein level develop a syndrome consisting of mitochondrial myopathy, cataract and combined respiratory chain deficiency (Di Fonzo, Ronchi et al. 2009, Calderwood, Holm et al. 2015, Nambot, Gavrilov et al. 2017). Our mechanistic studies demonstrated that *ALR* downregulation impairs mitochondrial import of ABCB8 protein, and decreased ABCB8 levels has been shown to cause defective cytosolic Fe/S cluster maturation (Ichikawa, Bayeva et al. 2012). This decreased mitochondrial ABCB8 level and ensured cytosolic Fe/S cluster maturation defect activates IRP1 and increases cellular iron uptake. Additionally, reduction in mitochondrial ABCB8 levels is also associated with increased sensitivity of cells to oxidative stress (Ichikawa, Bayeva et al. 2012). These two mechanisms may explain the profound muscle damage observed in patients carrying *ALR* mutations. It is also worth pointing out that elevated iron level has been causally linked to the cardiomyopathy phenotype in mice with cardiac-specific ABCB8 deletion (Chang, Wu et al. 2016). Therefore, it would be of great interest to determine genetic or pharmacologic modulation of cellular iron as a therapy for these patients.

Tissue samples from patients with ALR mutation demonstrated depressed mitochondrial respiratory chain activity (Di Fonzo, Ronchi et al. 2009, Calderwood, Holm et al. 2015). In contrast, our data (***Figure 1F and G***) and yeast studies (Lange, Lisowsky et al. 2001) demonstrated that ALR downregulation does not affect mitochondrial Fe/S cluster assembly, and similar results have been demonstrated with ABCB8 downregulation (Ichikawa, Bayeva et al. 2012). There could be two potential explanations for the discrepancy. First of all, not all reported patients with ALR mutations have complex IV deficiency. Therefore, there are likely other disease modifier genes explaining this incomplete penetrance of phenotype. Additionally, the difference between these findings may be due to acute versus chronic depletion of ALR. Chronic mitochondrial iron overload, a cellular response seen in many models of cytoplasmic Fe/S cluster maturation defects (Lange, Lisowsky et al. 2001, Pondarré, Antiochos et al. 2006, Ichikawa, Bayeva et al. 2012), has been linked to mitochondrial damage. Extensive mitochondrial damage has also been demonstrated in tissue samples from patients with *ALR* mutations (Nambot, Gavrilov et al. 2017). It is therefore not surprising that respiratory complex activities will also be reduced in cells full of defective mitochondria. Taken together, respiratory chain deficiency in those patients is likely to be a secondary effect of chronic cytoplasmic Fe/S cluster deficiency.

Mitochondrial proteins utilize different sets of importers to reach their final destination. Import of intermembrane space proteins requires cooperation between the TOM complex and the MIA40/ALR machinery. On the other hand, proper insertion of mitochondrial inner membrane proteins requires both the TOM complex and either the TIM22 or the TIM23 complex (Webb and Lithgow 2010). Both Atm1p, the yeast homolog of ABCB7, and Mdl1p, which shares significant sequence homology with ABCB8, utilize the Tim23 complex for mitochondrial import in a membrane potential dependent manner (Webb and Lithgow 2010, Stiller, Höpker et al. 2016). In this paper, we demonstrated that the transport of mammalian ABCB8 into the mitochondria is also TIM23-dependent. Additionally, we showed that mitochondrial import of ABCB8 is MIA40/ALR dependent. One potential explanation for the dual dependence of ABCB8 on MIA40/ALR protein import machinery as well as TIM23 complex for mitochondrial transport is that MIA40/ALR import machinery is important for the function of TIM23 complex. However, downregulation of MIA40 or ALR does not affect mitochondrial import of the canonical TIM23 complex substrate SU9-eGFP (***Figure 5-figure supplement 1C and D***), and the function of the Tim23 complex was not altered in yeast lacking Mia40 (Wrobel, Trojanowska et al. 2013). Therefore, modulation of MIA40/ALR complex likely affects the mitochondrial of ABCB8 import upstream of TIM23 and import into the inner mitochondrial membrane. We thus propose a model where a nascent ABCB8 peptide interacts with the MIA40/ALR import system and TIM23 complex sequentially during mitochondrial import, similar to the mechanism that is reported for p53 and Mrp10 for mitochondrial import (Zhuang, Wang et al. 2013, Longen, Woellhaf et al. 2014). Taken together, we have identified the MIA40/ALR system as a novel player in mitochondrial import of ABCB8 through direct interaction with the newly synthesized peptide, which occurs prior to peptide insertion into the inner mitochondrial membrane via the TIM23 complex (***Figure 6F***).

Our results demonstrate physical interaction between MIA40 and ABCB8 and suggest that ABCB8 is a novel substrate for the MIA40/ALR protein import system. Our sequence alignment identifies five cysteine residues that are preserved throughout higher vertebrates. These cysteines likely play a redundant function in the interaction between MIA40 and ABCB8, as the physical interaction between these ABCB8 and MIA40 did not diminish until all five cysteines were mutated. Furthermore, significantly less ABCB8 was copurified with MIA40 under a reducing condition, thereby supporting the model that ABCB8 interacts with MIA40 partially through disulfide bond formation. It is worth noting that the five conserved cysteines, or any other cysteine residues in ABCB8, do not form a CX2C or a CX9C consensus motif that is characteristic of most MIA40 substrates (Sideris, Petrakis et al. 2009). Therefore, how MIA40 recognize these residues and how this molecular interaction facilitates mitochondrial transport of ABCB8 remains to be determined.

Although our proposed model explains how ALR is involved in cellular iron homeostasis, this mechanism likely presented late in the evolution process. Our multi-species alignment of ABCB8 homologs revealed five conserved cysteines in higher vertebrates, and our experimental evidence argues that all five cysteines are important in the interaction with MIA40. However, not all five cysteines are conserved in lower vertebrates such as zebrafish and xenopus. Additionally, the functional homolog of ABCB8 in yeast has not been identified yet. While it was speculated that Mdl1p was the yeast homolog of mammalian ABCB8 (Ichikawa, Bayeva et al. 2012), Mdl1p only shared 28% sequence identity and additional 34% sequence similarity with human ABCB8 protein. Most of the sequence similarities reside in the ATPase domain of the protein. Therefore, our proposed mechanism is likely a late product of evolution and a similar mechanism may not exist in yeast.

In summary, we demonstrated that cytosolic Fe/S cluster maturation in mammalian cells depends on the mitochondrial isoform of ALR, which mediates the transport of ABCB8 into mitochondria. ALR thus plays a role in cellular iron regulation through indirectly regulating IRP1 activity. Sequence variation in conserved cysteine residues in ABCB7 and ABCB8 may explain why the mitochondrial import of only ABCB8 but not ABCB7 depends on the MIA40/ALR pathway. This report presents the first mechanism of how seemingly unrelated mitochondrial proteins involved in sulfur redox homeostasis and mitochondrial protein import are linked together in the cytosolic Fe/S cluster maturation pathway, and provides additional insights into cellular iron regulation, with possible implications in diseases presenting with altered cellular iron homeostasis and Fe/S cluster maturation.

## Materials and methods

### Cloning

*Tfrc* promoter reporter construct was generated by cloning a fragment from −897bp to +129bp around mouse TfR1 transcription site into pGL3basic reporter. *Tfrc* 3’UTR luciferase construct was described previously (Bayeva, Khechaduri et al. 2012). IRE sites in this construct were deleted using QuickChange Site-Directed Mutagenesis Kit (Agilent). pLenti-rtTA3, SU9-eGFP and pBI-eGFP plasmids were purchased from Addgene. Human ABCB7 cDNA plasmid was a generous gift from Dr. Berry Paw (Brigham and Women’s Hospital and Boston Children’s Hospital). Human MIA40 and ALR cDNA clones were purchased from Open BioSystems. Coding sequence of Mia40, ALR, ABCB8, SU9-eGFP, and ABCB7weresubcloned into pBI-eGFP for import experiment. ABCB8 with 6X-His or 3X-Flag tag in the C terminus and Mia40 with 6X-His tag in the C terminus were inserted into pCMV6-XL6. Cysteine mutants of ABCB8 were generated using PCR-based site-directed mutagenesis. All constructs were sequenced before experiments.

### Cell culture

*Ireb2* KO MEFs and wild type MEFs isolated from littermate control were generous gifts from Dr. Tracy Rouault (NICHD, NIH). Mouse embryonic fibroblasts were cultured in DMEM (Cellgro) supplemented with 10% FBS, 100U/ml penicillin and 100U/ml streptomycin. HEK293 cells were cultured in MEM (Cellgro) supplemented with 10% FBS, 100U/ml penicillin, 100U/ml streptomycin, and 1mM sodium pyruvate.

### Lentivirus production

Full length and cytoplasmic isoform of ALR were cloned into pHIV-eGFP vector. The resultant vector was packaged into lentiviral vector in HEK293T cells after cotransfection with pSPAX and pMD2.G plasmids. The culture supernatant was mixed with polybrene to final concentration of 15μg/ml and used to infect MEFs for overexpressing isoform-specific ALR. pLenti-rtTA3 was packaged into lentiviral vector in HEK293T cells after cotransfection with pSPAX and pMD2.G plasmids. Culture supernatant was used to infect HEK293 as described above, and cells with stable integration of viral genome were selected with Blasticidin (Invitrogen).

### Transfection

siRNA against human *ALR, MIA40, TIM23*, and mouse *Alr* and *Aco1* were purchased from Dharmacon. siRNA against 3’UTR sequence of mouse *Alr* were purchased from Qiagen. siRNAs were transfected into HEK293 or MEF using Dharmafect 1 Transfection Reagent (Dharmacon). Plasmids were transfected into HEK293 cells using either Lipofectamine 2000 reagent (Invitrogen) or calcium phosphate transfection.

### Reverse Transcription and Quantitative Realtime PCR

RNA was isolated from cells using RNA-STAT60 (Tel-Test), and reverse transcribed with Taqman Reverse Transcription Reagents (Invitrogen) according to manufacturers’ instruction. Quantification of relative gene expression was done using Fast SYBR Green Master Mix (Applied Biosystems) and run on 7500 Fast Real-Time PCR system (Applied Biosystems).

### Cellular Uptake of ^55^Fe-transferrin and steady state cellular iron content

^55^FeCl_3_ (Perkin Elmer) was first conjugated to nitrilotriacetic acid (NTA, Sigma) before adding transferrin (Tf) in 2:1 molar ratio. The mix was incubated at room temperature for one hour. ^55^Fe-transferrin was separated from unbound ^55^Fe by running on PD-10 desalting column (GE Healthcare). For uptake experiment, cells were incubated with media containing 55μg/ml ^55^Fe-Tf for two hours. Residual membrane-bound iron was removed by washing cells with cold 200μM DFO in PBS. Radioactivity was quantified on a Beckman scintillation counter, and the results were normalized to cellular protein content of the same sample. For steady state cellular ^55^Fe content, cells were incubated with 280μM of ^55^FeNTA for 2 days. The subcellular fractions were isolated and radioactivity in the fraction was measured as described above.

### Mitochondrial Fractioning

Mitochondria from cells were purified using Mitochondrial Isolation Kit for Cells (Pierce) according to manufacturer’s instruction.

### Enzyme activity assay

Complex I and II activity were measured with Complex I and Complex II Enzyme Activity Microplate Assay Kit (MitoScience), respectively, according to manufacturer’s instruction. Xanthine oxidase activity was measured using AmplexRed Xanthine Oxidase Activity Kit (Invitrogen). For cytosolic aconitase assay, cytosolic fraction was concentrated on Amicon Ultra-0.5 Centrifugal Filter Unit with 10kDa cutoff (Millipore) and the buffer was exchanged to PBS. The aconitase activity in the concentrated fraction was measured using Aconitase Activity Assay kit (MitoScience).

### Luciferase Assay

Twenty-four hours after transfection with Firefly reporter construct and Renilla luciferase construct (as normalization control for transfection efficiency), cells were solubilized in 1X passive lysis buffer (Promega). The lysates were loaded onto a 96 well plate, and the luciferase activity was determined using Dual Luciferase Assay System (Promega) with Modulus Microplate Reader (Turner Biosystems). The ratio between firefly and renilla luciferase activities was normalized to that of empty vector under corresponding treatments.

### Western Blotting

Proteins were resolved on 4-12% Novex Bis-Tris poly-acrylamide gel (Invitrogen) and blotted onto nitrocellulose membrane (Invitrogen). Membranes were incubated with primary antibodies for ALR (ProteinTech), GAPDH and α-tubulin (Abcam), GPAT (a generous gift from Dr. Roland Lill (University of Marburg)), or 70kDa subunit SDH (Invitrogen) for overnight, before the addition of HRP-conjugated secondary antibodies (Santa Cruz). The presence of target protein was visualized using Super Surgical Western Pico ECL substrate (Pierce). Quantification of western blotting image was done using ImageJ (NIH).

### Mitochondrial import of target protein

36 hours after transfection, cells were treated with 1ug/ml doxycycline, and harvested after 6 hours for mitochondrial fractionation. For CCCP treatment, 2.5uM CCCP was added at the 4^th^ hour and the cells were incubated for additional 2 hours. The mitochondrial level of protein of interest is normalized to the GFP levels in the cytosolic fraction from the same sample.

### Ni-NTA pulldown assay

Mitochondrial pallets were lysed in PBS containing 1% Triton X100 (Sigma), protease inhibitor (G Biosciences) and 10mM immidazole (GE Healthcare). The lysate was cleared via centrifugation and loaded to His Spin Trap spin column (GE Healthcare). The column was sequentially washed with PBC containing 20mM, 40mM, and 80mM immidazole and eluted with PBS containing 200mM immidazole. 100mM DTT was included in the lysis buffer when indicated.

### Statistical Analysis

Data are presented as mean ± SEM. Statistical significance was assessed with ANOVA, with post hoc Tukey’s test for multiple group comparison. A P-value less than 0.05 was considered statistically significant.

## Acknowledgements

We would like to thank Drs. Roland Lill, Barry Paw, Tracy Rouault, and Celeste Simon for sharing reagents. We would also like to thank technical help from Chunlei Chen and Justin Geier. H.-C.C. is supported by American Heart Association 12PRE12030002 and NIH T32 GM008152. H.A. is supported by NIH R01 HL127646, R01 HL140973, and R01 HL138982.

## Author Contributions

H.-C.C. and H.A. designed and performed the research and wrote the manuscript. J.S.S., X.J., T.S., and G.S. performed experiments. K.T.S. and J.S.S. provided critical reviews and feedbacks on the manuscript. H.A. supervised the project.

---

**Figure 2-figure supplement 1.**
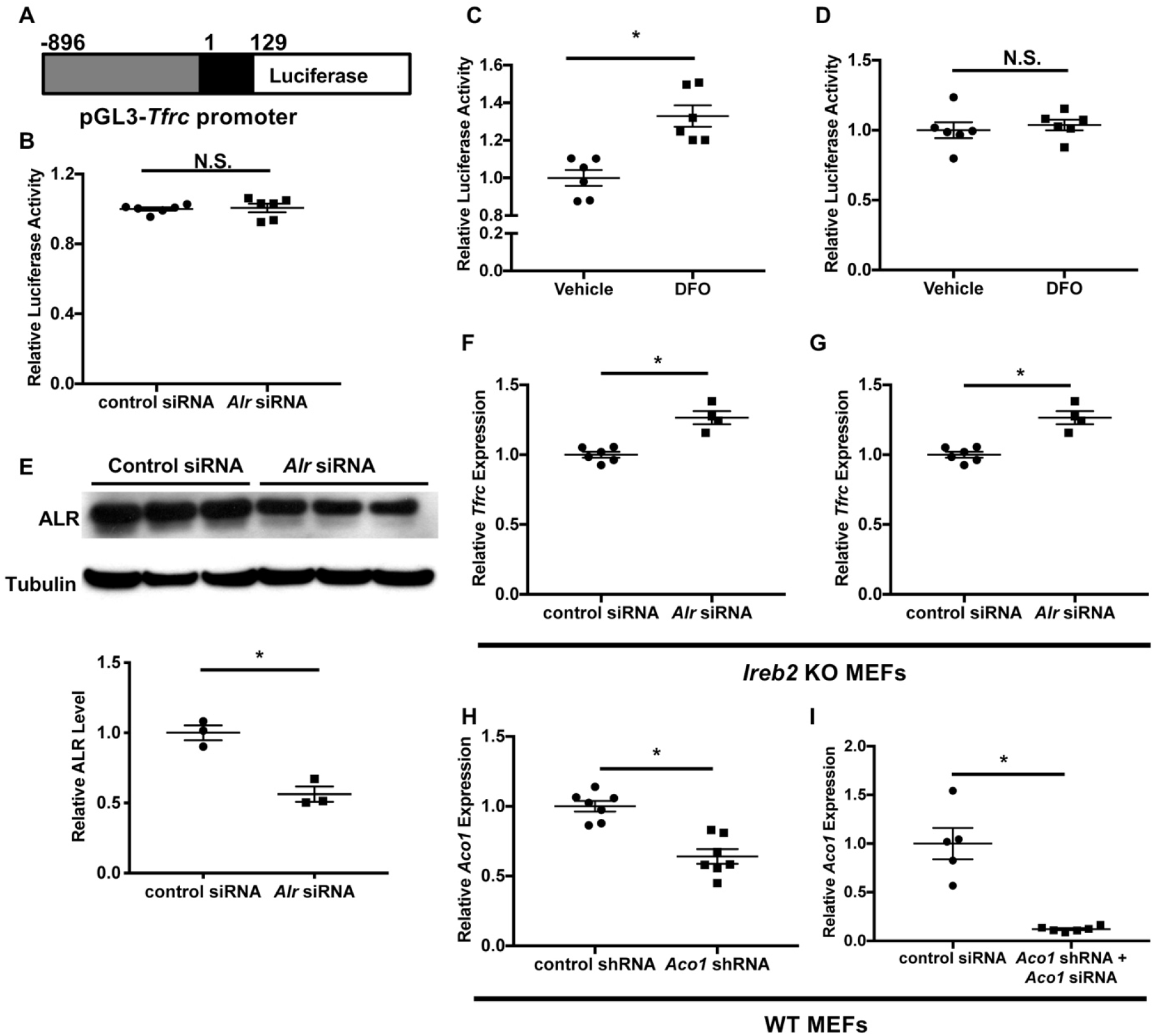
ALR regulates *Tfrc* mRNA levels through a post-transcriptional mechanism independent of IRP2. (**A**) Schematic representation of the *Tfrc* promoter reporter construct. +1 represent transcription start site on *Tfrc* genomic sequence. (**B**) *Tfrc* promoter activities are measured in wildtype MEFs with *Alr* downregulation (n=6). Luficerase activity of full length Tfrc 3’UTR (n=6, **C**) and Tfrc 3’UTR ΔIRE (n=6, **D**) constructs in cells treated with vehicle or DFO. (**E**) Alr protein levels in *Ireb2* KO MEFs treated with *Alr* siRNA (n=3). Densitometry analysis is shown below the western blots. (**F**) *Tfrc* mRNA level in *Ireb2* KO MEFs treated with *Alr* siRNA (n=4-6). (**G**) ^55^Fe-transferrin uptake in *Irrb2* KO MEFs treated with *Alr* siRNA (n=6). (**H**) *Aco1* mRNA levels in WT MEFs stably expressing *Aco1* shRNA at baseline (n=6). (**I**) *Aco1* mRNA levels in WT MEFs stably expressing *Aco1* shRNA and transfected with *Aco1* siRNA (n=5-6). Data are presented as mean ± SEM.* P<0.05 by ANOVA. N.S. = not significant.

**Figure 3-figure supplement 1.**
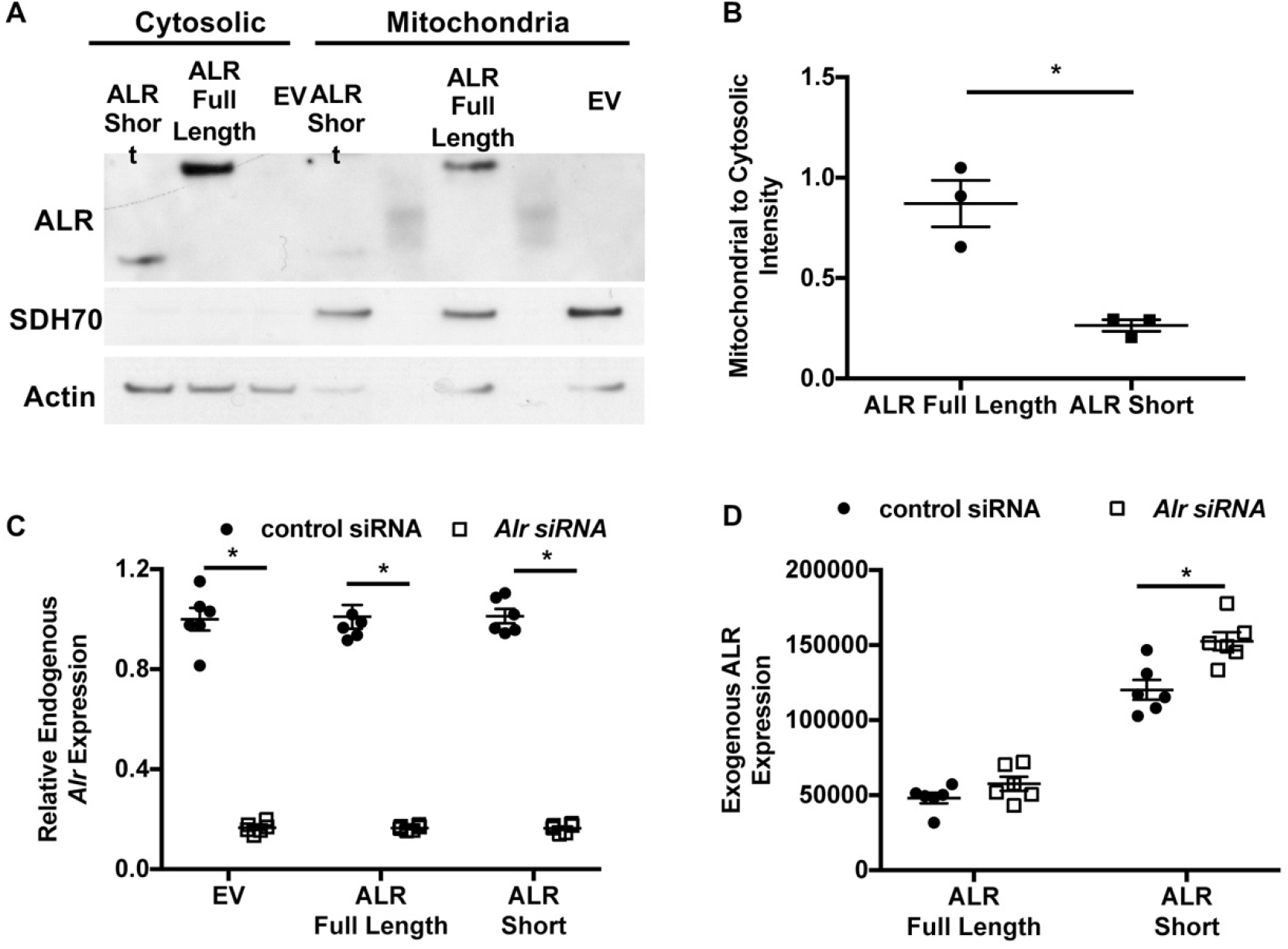
Validation of isoform-specific overexpression of ALR with concurrent downregulation of endogenous *Alr*. (**A**) Representative western blotting results showing the subcellular localization of full length and cytosolic isoform of ALR. (**B**) Densitometry analysis mitochondrial localization of each construct shown in panel **A** (n=3). (**C**) Expression of endogenous *Alr* in WT MEFs treated with *Alr* siRNA targeting 3’UTR of the endogenous locus and overexpressing indicated constructs (n=6). (**D**) mRNA levels of overexpressed ALR in WT MEFs treated with *Alr* siRNA targeting 3’UTR of the endogenous locus and overexpressing indicated constructs (n=6). EV = empty vector. ALR short = truncated ALR lacking the first 80 amino acids. Data are presented as mean ± SEM.* P<0.05 by ANOVA.

**Figure 4-figure supplement 1.**
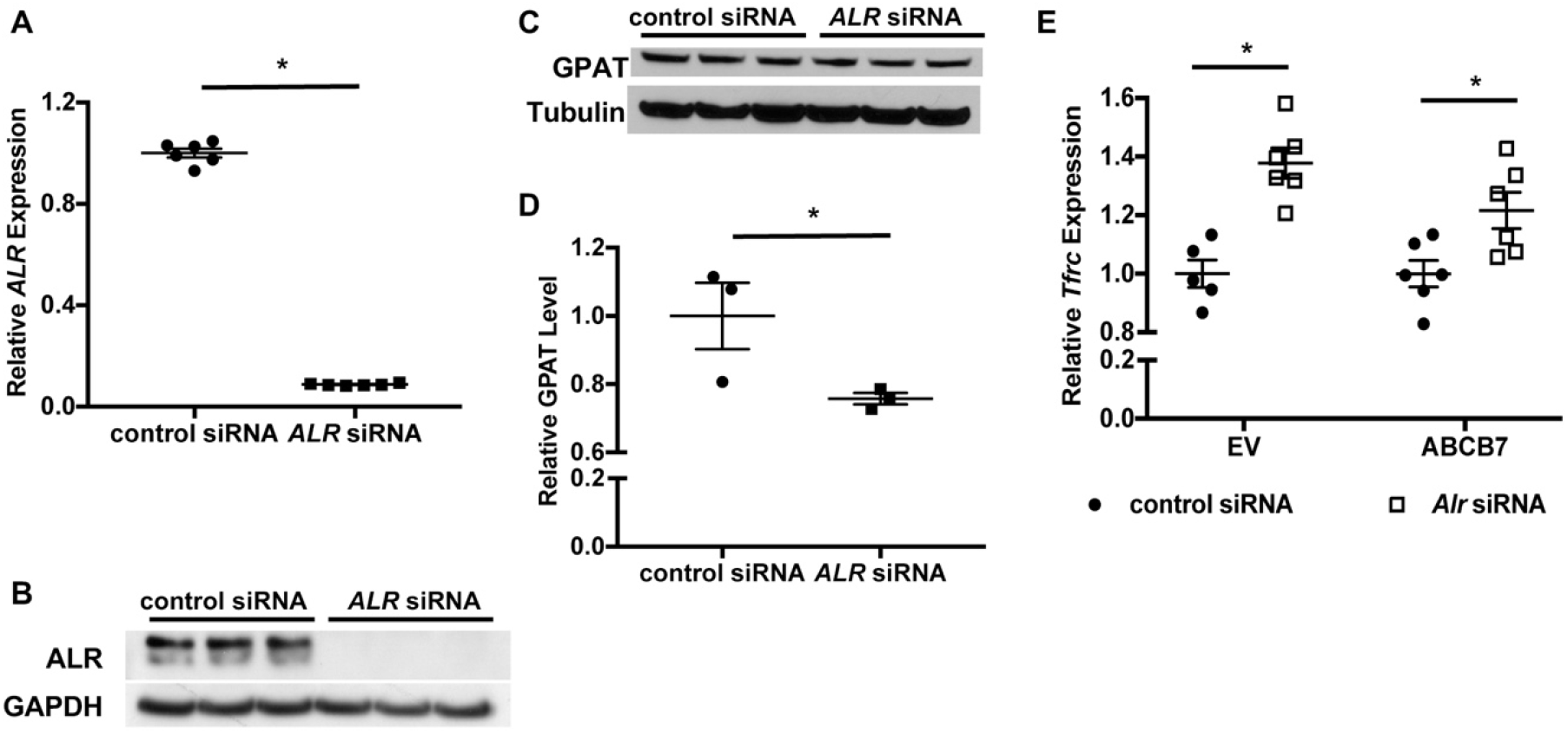
Downregulation of *ALR* in HEK293 cells results in cytosolic Fe/S cluster maturation defects. (**A**) *ALR* mRNA levels in HEK293 cells treated with *ALR* siRNA (n=6). (**B**) Representative western blot showing ALR protein levels in HEK293 cells treated with *ALR* siRNA (n=3). (**C**) Representative western blot showing GPAT protein levels in HEK293 cells treated with *ALR* siRNA (n=3). (**D**) Densitometry analysis of western blots shown in in panel **C**. (**E**) *Tfrc* mRNA levels in WT MEFs overexpressing ABCB7 and/or treated with *Alr* siRNA (n=5-6). Data are presented as mean ± SEM.* P<0.05 by ANOVA.

**Figure 5-figure supplement 1.**
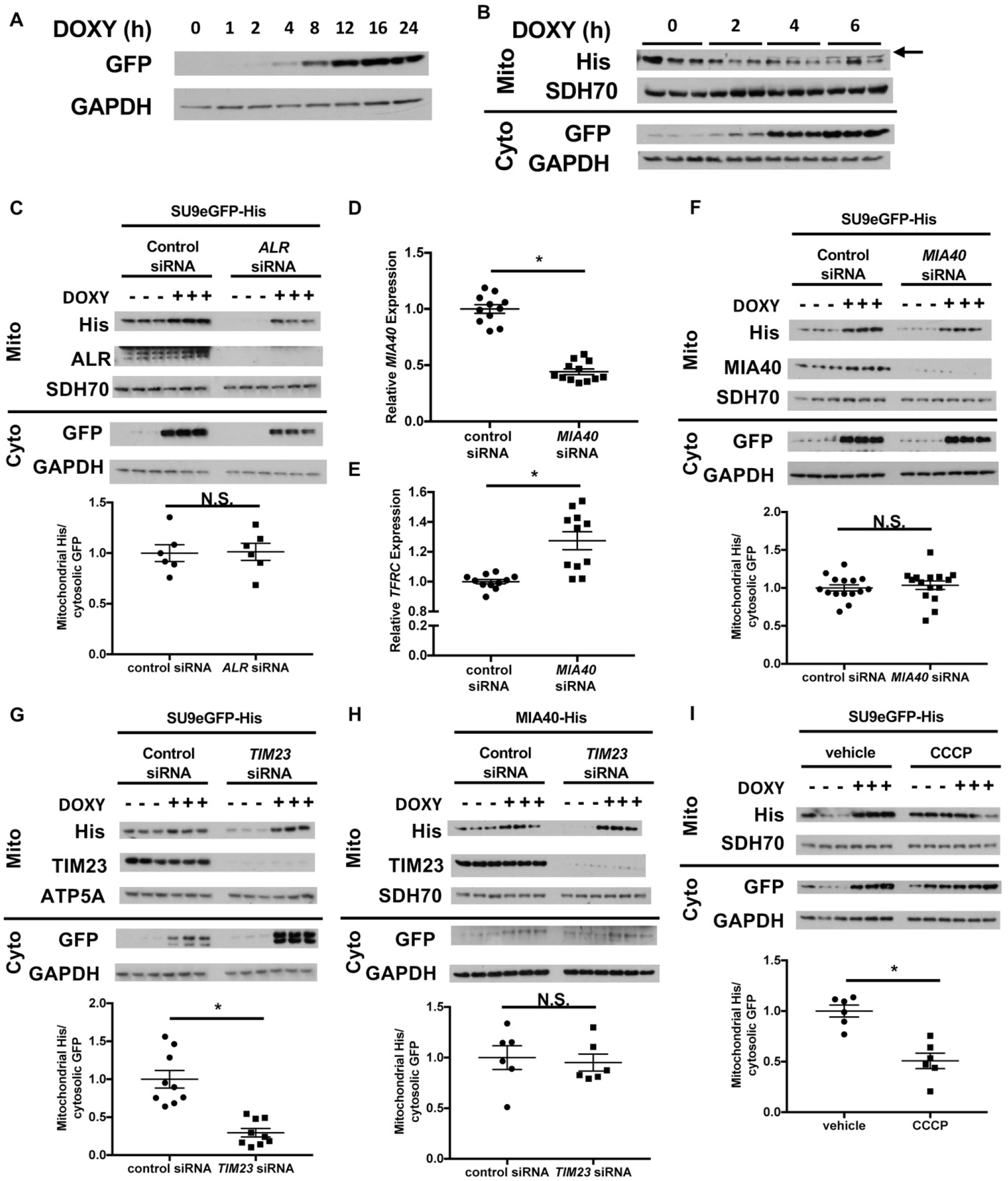
Downregulation of *ALR* or *MIA40* does not affect mitochondrial import of SU9-eGFP. (**A**) Representative western blot showing time dependent accumulation of GFP protein in cells stably expressing rtTA3 and transfected with doxycycline-inducible GFP expression plasmid. (**B**) Representative western blot images showing time dependent accumulation of GFP in the cytosol and ABCB7 in the mitochondria in cells stably expressing rtTA3 and transfected with the bicistronic doxycycline-inducible plasmid containing GFP and ABCB7. Arrow indicates the correct band for ABCB7. (**C**) Imported levels of SU9eGFP-His in cells with or without *ALR* downregulation with and without doxycycline treatment (n=6). *MIA40* (**D**) and *TFRC* (**E**) mRNA levels in HEK293 cells treated with *MIA40* siRNA (n=11-12). (**F**) Imported levels of SU9eGFP-His in cells with or without *MIA40* downregulation with and without doxycycline treatment (n=15). (**G**) Imported levels of SU9eGFP-His in cells with or without *TIM23* downregulation with and without doxycycline treatment (n=9). (**H**) Imported levels of MIA40-His in cells with or without *TIM23* downregulation with and without doxycycline treatment (n=6). (**I**) Imported levels of SU9eGFP-His in cells treated with CCCP and/ or doxycycline treatment (n=6). Mito = mitochondrial fraction. Cyto = cytosolic fraction. DOXY = doxycycline. Data are presented as mean ± SEM.* P<0.05 by ANOVA. N.S. = not significant.

**Figure 6-figure supplement 1.**
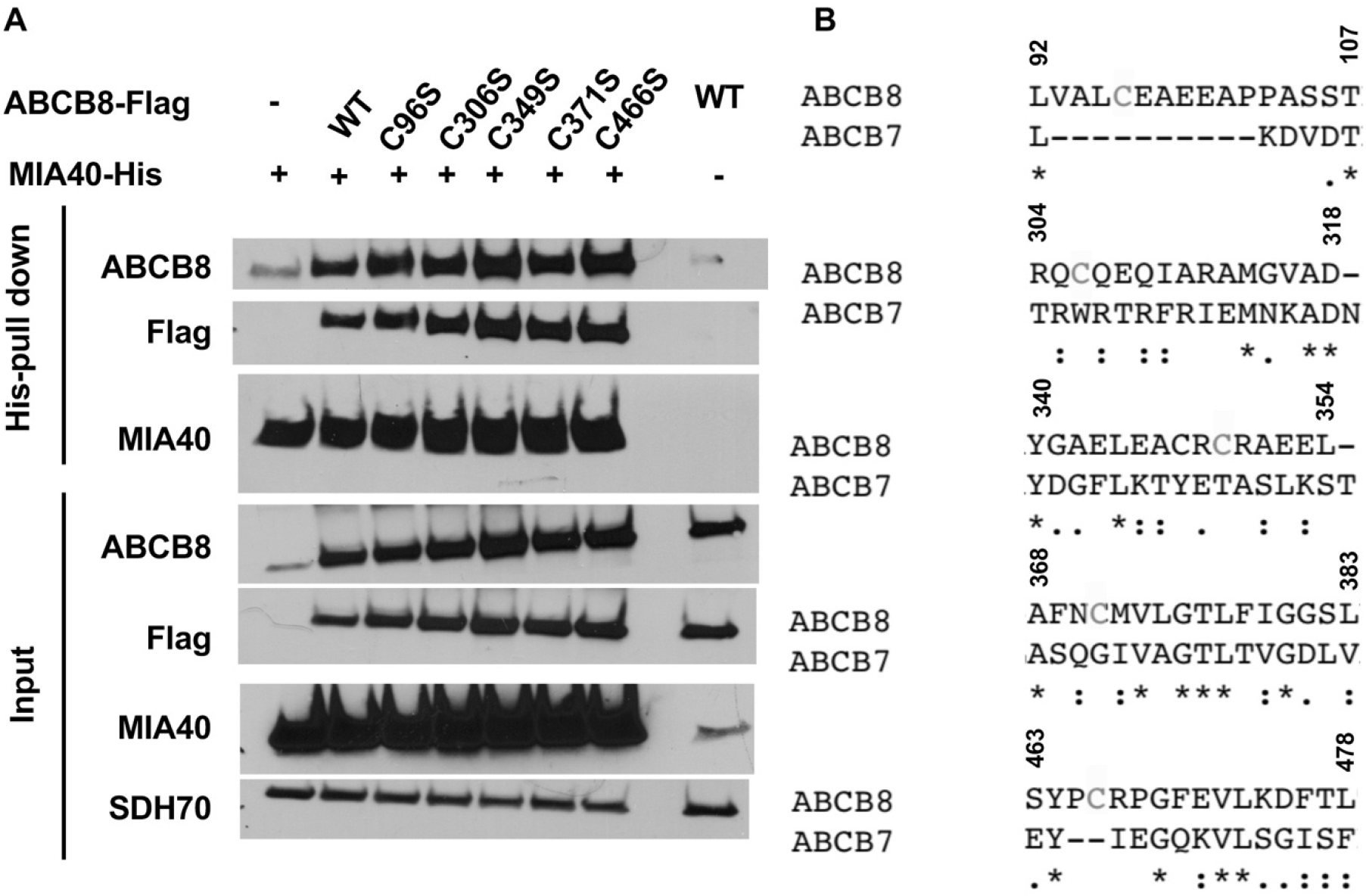
Individual mutation of the five conserved cysteines in ABCB8 minimally impacts its interaction with MIA40. (**A**) Western blot results showing co-immunoprecipitation of His-tagged MIA40 and various mutants of Flag-tagged ABCB8 in cells overexpressing indicated constructs. (**B**) Clustal Omega alignment between human ABCB7 and ABCB8 protein sequences demonstrate lack of conservation of the cysteines.

## References

Anderson, C. P., M. Shen, R. S. Eisenstein and E. A. Leibold (2012). “Mammalian iron metabolism and its control by iron regulatory proteins.” Biochimica et Biophysica Acta (BBA) - Molecular Cell Research 1823(9): 1468–1483.

Banci, L., I. Bertini, V. Calderone, C. Cefaro, S. Ciofi-Baffoni, A. Gallo, E. Kallergi, E. Lionaki, C. Pozidis and K. Tokatlidis (2011). “Molecular recognition and substrate mimicry drive the electron-transfer process between MIA40 and ALR.” Proceedings of the National Academy of Sciences 108(12): 4811–4816.

Bayeva, M., H.-C. Chang, R. Wu and H. Ardehali (2013). “When less is more: novel mechanisms of iron conservation.” Trends in Endocrinology & Metabolism 24(11): 569–577.

Bayeva, M., A. Khechaduri, S. Puig, H.-C. Chang, S. Patial, Perry J. Blackshear and H. Ardehali (2012). “mTOR Regulates Cellular Iron Homeostasis through Tristetraprolin.” Cell Metabolism 16(5): 645–657.

Calderwood, L., I. A. Holm, L. A. Teot and I. Anselm (2015). “Adrenal Insufficiency in Mitochondrial Disease: A Rare Case of GFER-Related Mitochondrial Encephalomyopathy and Review of the Literature.” Journal of Child Neurology 31(2): 190–194.

Chang, H.-C., R. Wu, M. Shang, T. Sato, C. Chen, J. S. Shapiro, T. Liu, A. Thakur, K. T. Sawicki, S. V. Prasad and H. Ardehali (2016). “Reduction in mitochondrial iron alleviates cardiac damage during injury.” EMBO Molecular Medicine 8(3): 247–267.

Dabir, Deepa V., Samuel A. Hasson, K. Setoguchi, Meghan E. Johnson, P. Wongkongkathep, Colin J. Douglas, J. Zimmerman, R. Damoiseaux, Michael A. Teitell and Carla M. Koehler (2013). “A Small Molecule Inhibitor of Redox-Regulated Protein Translocation into Mitochondria.” Developmental Cell 25(1): 81–92.

Daithankar, V. N., S. A. Schaefer, M. Dong, B. J. Bahnson and C. Thorpe (2010). “Structure of the Human Sulfhydryl Oxidase Augmenter of Liver Regeneration and Characterization of a Human Mutation Causing an Autosomal Recessive Myopathy.” Biochemistry 49(31): 6737–6745.

De Domenico, I., D. McVey Ward and J. Kaplan (2008). “Regulation of iron acquisition and storage: consequences for iron-linked disorders.” Nat Rev Mol Cell Biol 9(1): 72–81.

Di Fonzo, A., D. Ronchi, T. Lodi, E. Fassone, M. Tigano, C. Lamperti, S. Corti, A. Bordoni, F. Fortunato, M. Nizzardo, L. Napoli, C. Donadoni, S. Salani, F. Saladino, M. Moggio, N. Bresolin, I. Ferrero and G. P. Comi (2009). “The Mitochondrial Disulfide Relay System Protein GFER Is Mutated in Autosomal-Recessive Myopathy with Cataract and Combined Respiratory-Chain Deficiency.” American journal of human genetics 84(5): 594–604.

Gandhi, C. R., J. R. Chaillet, M. A. Nalesnik, S. Kumar, A. Dangi, A. J. Demetris, R. Ferrell, T. Wu, S. Divanovic, T. Stankeiwicz, B. Shaffer, D. B. Stolz, S. A. Harvey, J. Wang and T. E. Starzl (2015). “Liver-specific deletion of augmenter of liver regeneration accelerates development of steatohepatitis and hepatocellular carcinoma in mice.” Gastroenterology 148(2): 379–391.e374.

Haindrich, A. C., M. Boudova, M. Vancova, P. P. Diaz, E. Horakova and J. Lukes (2017). “The intermembrane space protein Erv1 of Trypanosoma brucei is essential for mitochondrial Fe-S cluster assembly and operates alone.” Mol Biochem Parasitol 214: 47–51.

Hentze, M. W., M. U. Muckenthaler, B. Galy and C. Camaschella (2010). “Two to Tango: Regulation of Mammalian Iron Metabolism.” Cell 142(1): 24–38.

Huang, L. L., R. T. Long, G. P. Jiang, X. Jiang, H. Sun, H. Guo and X. H. Liao (2018). “Augmenter of liver regeneration promotes mitochondrial biogenesis in renal ischemia-reperfusion injury.” Apoptosis 23(11-12): 695–706.

Ichikawa, Y., M. Bayeva, M. Ghanefar, V. Potini, L. Sun, R. K. Mutharasan, R. Wu, A. Khechaduri, T. Jairaj Naik and H. Ardehali (2012). “Disruption of ATP-binding cassette B8 in mice leads to cardiomyopathy through a decrease in mitochondrial iron export.” Proceedings of the National Academy of Sciences.

Jensen, R. E. and C. D. Dunn (2002). “Protein import into and across the mitochondrial inner membrane: role of the TIM23 and TIM22 translocons.” Biochimica et Biophysica Acta (BBA) - Molecular Cell Research 1592(1): 25–34.

Kallergi, E., M. Andreadaki, P. Kritsiligkou, N. Katrakili, C. Pozidis, K. Tokatlidis, L. Banci, I. Bertini, C. Cefaro, S. Ciofi-Baffoni, K. Gajda and R. Peruzzini (2012). “Targeting and Maturation of Erv1/ALR in the Mitochondrial Intermembrane Space.” ACS Chemical Biology 7(4): 707–714.

Kumar, S., R. Rani, R. Karns and C. R. Gandhi (2019). “Augmenter of liver regeneration protein deficiency promotes hepatic steatosis by inducing oxidative stress and microRNA-540 expression.” Faseb j 33(3): 3825–3840.

Kumar, S., J. Wang, R. Rani and C. R. Gandhi (2016). “Hepatic Deficiency of Augmenter of Liver Regeneration Exacerbates Alcohol-Induced Liver Injury and Promotes Fibrosis in Mice.” PLoS One 11(1): e0147864.

Lange, H., T. Lisowsky, J. Gerber, U. Muhlenhoff, G. Kispal and R. Lill (2001). “An essential function of the mitochondrial sulfhydryl oxidase Erv1p/ALR in the maturation of cytosolic Fe/S proteins.” EMBO Rep 2(8): 715–720.

Li, Y., K. Wei, C. Lu, Y. Li, M. Li, G. Xing, H. Wei, Q. Wang, J. Chen, C. Wu, H. Chen, S. Yang and F. He (2002). “Identification of hepatopoietin dimerization, its interacting regions and alternative splicing of its transcription.” European Journal of Biochemistry 269(16): 3888–3893.

Lill, R., R. Dutkiewicz, H.-P. Elsässer, A. Hausmann, D. J. A. Netz, A. J. Pierik, O. Stehling, E. Urzica and U. Mühlenhoff (2006). “Mechanisms of iron–sulfur protein maturation in mitochondria, cytosol and nucleus of eukaryotes.” Biochimica et Biophysica Acta (BBA) - Molecular Cell Research 1763(7): 652–667.

Liu, L., P. Xie, W. Li, Y. Wu and W. An (2019). “Augmenter of Liver Regeneration Protects against Ethanol-Induced Acute Liver Injury by Promoting Autophagy.” The American Journal of Pathology 189(3): 552–567.

Longen, S., Michael W. Woellhaf, C. Petrungaro, J. Riemer and Johannes M. Herrmann (2014). “The Disulfide Relay of the Intermembrane Space Oxidizes the Ribosomal Subunit Mrp10 on Its Transit into the Mitochondrial Matrix.” Developmental Cell 28(1): 30–42.

Müller, J. M., D. Milenkovic, B. Guiard, N. Pfanner and A. Chacinska (2008). “Precursor Oxidation by Mia40 and Erv1 Promotes Vectorial Transport of Proteins into the Mitochondrial Intermembrane Space.” Molecular Biology of the Cell 19(1): 226–236.

Nambot, S., D. Gavrilov, J. Thevenon, A. L. Bruel, M. Bainbridge, M. Rio, C. Goizet, A. Rötig, J. Jaeken, N. Niu, F. Xia, A. Vital, N. Houcinat, F. Mochel, P. Kuentz, D. Lehalle, Y. Duffourd, J. B. Rivière, C. Thauvin-Robinet, A. L. Beaudet and L. Faivre (2017). “Further delineation of a rare recessive encephalomyopathy linked to mutations in GFER thanks to data sharing of whole exome sequencing data.” Clinical Genetics 92(2): 188–198.

Peleh, V., E. Cordat and J. M. Herrmann (2016). “Mia40 is a trans-site receptor that drives protein import into the mitochondrial intermembrane space by hydrophobic substrate binding.” eLife 5: e16177.

Polimeno, L., B. Pesetti, E. Annoscia, F. Giorgio, R. Francavilla, T. Lisowsky, A. Gentile, R. Rossi, A. Bucci and A. Francavilla (2011). “Alrp, a survival factor that controls the apoptotic process of regenerating liver after partial hepatectomy in rats.” Free Radical Research 45(5): 534–549.

Pondarré, C., B. B. Antiochos, D. R. Campagna, S. L. Clarke, E. L. Greer, K. M. Deck, A. McDonald, A.-P. Han, A. Medlock, J. L. Kutok, S. A. Anderson, R. S. Eisenstein and M. D. Fleming (2006). “The mitochondrial ATP-binding cassette transporter Abcb7 is essential in mice and participates in cytosolic iron–sulfur cluster biogenesis.” Human Molecular Genetics 15(6): 953–964.

Sato, K., Y. Torimoto, T. Hosoki, K. Ikuta, H. Takahashi, M. Yamamoto, S. Ito, N. Okamura, K. Ichiki, H. Tanaka, M. Shindo, K. Hirai, Y. Mizukami, T. Otake, M. Fujiya, K. Sasaki and Y. Kohgo (2011). “Loss of ABCB7 gene: pathogenesis of mitochondrial iron accumulation in erythroblasts in refractory anemia with ringed sideroblast with isodicentric (X)(q13).” Int J Hematol 93(3): 311–318.

Sideris, D. P., N. Petrakis, N. Katrakili, D. Mikropoulou, A. Gallo, S. Ciofi-Baffoni, L. Banci, I. Bertini and K. Tokatlidis (2009). “A novel intermembrane space-targeting signal docks cysteines onto Mia40 during mitochondrial oxidative folding.” J Cell Biol 187(7): 1007–1022.

Song, D. and F. S. Lee (2011). “Mouse knock-out of IOP1 protein reveals its essential role in mammalian cytosolic iron-sulfur protein biogenesis.” J Biol Chem 286(18): 15797–15805.

Stiller, Sebastian B., J. Höpker, S. Oeljeklaus, C. Schütze, Sandra G. Schrempp, J. Vent-Schmidt, Susanne E. Horvath, Ann E. Frazier, N. Gebert, M. van der Laan, M. Bohnert, B. Warscheid, N. Pfanner and N. Wiedemann (2016). “Mitochondrial OXA Translocase Plays a Major Role in Biogenesis of Inner-Membrane Proteins.” Cell Metabolism 23(5): 901–908.

Sztolsztener, M. E., A. Brewinska, B. Guiard and A. Chacinska (2013). “Disulfide Bond Formation: Sulfhydryl Oxidase ALR Controls Mitochondrial Biogenesis of Human MIA40.” Traffic 14(3): 309–320.

Tacchini, L., E. Gammella, C. De Ponti, S. Recalcati and G. Cairo (2008). “Role of HIF-1 and NF-κB Transcription Factors in the Modulation of Transferrin Receptor by Inflammatory and Antiinflammatory Signals.” Journal of Biological Chemistry 283(30): 20674–20686.

Todd, L. R., R. Gomathinayagam and U. Sankar (2010). “A novel Gfer-Drp1 link in preserving mitochondrial dynamics and function in pluripotent stem cells.” Autophagy 6(6): 821–822.

Webb, C. T. and T. Lithgow (2010). “Mitochondrial Biogenesis: Sorting Mechanisms Cooperate in ABC Transporter Assembly.” Current Biology 20(13): R564–R567.

Wrobel, L., A. Trojanowska, M. E. Sztolsztener and A. Chacinska (2013). “Mitochondrial protein import: Mia40 facilitates Tim22 translocation into the inner membrane of mitochondria.” Mol Biol Cell 24(5): 543–554.

Ye, H. and T. A. Rouault (2010). “Human Iron-Sulfur Cluster Assembly, Cellular Iron Homeostasis, and Disease.” Biochemistry 49(24): 4945–4956.

Zhuang, J., P.-y. Wang, X. Huang, X. Chen, J.-G. Kang and P. M. Hwang (2013). “Mitochondrial disulfide relay mediates translocation of p53 and partitions its subcellular activity.” Proceedings of the National Academy of Sciences 110(43): 17356–17361.

